# In-silico Tandem Mass Spectral Libraries for the Blood Exposome and Metabolomics Workbench Databases: Application of Gemini Large Language Model

**DOI:** 10.64898/2026.10.20.760278

**Authors:** Tuan Nguyen, Dinesh Barupal

**Affiliations:** Integrated Data Science Laboratory for Metabolomics and Exposomics, Department of Environmental Medicine, Icahn School of Medicine at Mount Sinai, New York, NY, 10029, USA

## Abstract

Human body contains endogenous metabolites, nutrients and xenobiotics chemicals. These chemicals can be detected in a biospecimen using high-resolution mass spectrometry instruments, yielding metabolomic and exposomic datasets. Mass spectral libraries are critical components of these workflows to annotate experimental data with chemical information. In this data descriptor, we report mass spectral libraries for human-relevant small molecules, that are generated using Google Gemini large-language model. Using an optimized chain-of-thought prompt and instructions, we have generated mass spectra for each compound at high/medium/low energy fragmentation level. Each spectrum was reported in the NIST MSP text format for easy to use by metabolomics software. These libraries cover 91,231 unique 2D structures from the Blood Exposome Database and the Metabolomics Workbench data repository. Technical validation was shown by comparing the Gemini-generated spectra against NIST 23 mass spectral database, which demonstrated 55% accuracy. LLMs represent a useful and accessible tool for expanding *in-silico* spectral libraries to annotate compounds in the HRMS data for human biospecimens and may accelerate the compound annotation in metabolomics and exposomics. Both mass spectral libraries are available at https://zenodo.org/records/20102592.

## Background and Summary

The human body represents a highly complex and dynamic chemical environment, comprising tens of thousands of endogenous metabolites and exogenous chemicals. These molecules originate from internal biochemical reactions as well as external sources such as environmental exposures, diet, and lifestyle factors^1,2^. Understanding this vast chemical space is crucial for unraveling fundamental biological processes and determining their clinical significance in disease pathology and health maintenance. The fields of metabolomics and exposomics aim to comprehensively profile these chemical signatures. Currently, major knowledgebases such as the Blood Exposome Database^1^, Metabolomics Workbench^3^, and KEGG^4^ catalog thousands of known compounds. These reference datasets are continually updated through rigorous literature mining, database fusion, and experimental validation. However, the biological function, origin, and spectral signatures of a vast portion of this chemical space remain unknown, representing the dark matter of the human metabolome and exposome.

Processing the massive amounts of data generated by high-resolution mass spectrometry (HRMS) relies primarily on matching experimental data against mass spectral libraries. Software tools such as IDSL.CSA^5^, MS-DIAL^6^, and the NIST MS Search program utilize spectral similarity scoring algorithms, such as cosine similarity and entropy similarity, to match experimental MS/MS spectra against reference libraries. The confidence levels of these annotations depend significantly on the library used. In-house libraries built with pure reference standards provide the highest confidence but are often limited in size due to the prohibitive cost and lack of commercial availability of many chemical standards. Public and commercial databases like NIST23 offer broader coverage but still capture only a fraction of the known chemical space. Consequently, a vast majority of detected features in untargeted metabolomics and exposomics remain unannotated^7,8^. To overcome the bottleneck of limited experimental reference spectra, researchers are increasingly using *in-silico* methods to predict mass spectra and expand these crucial libraries^9–12^.

A majority of compounds in the metabolomic and exposomic knowledgebases lack mass spectral data^10^, limiting their use in annotating high-resolution mass spectrometry datasets. Collecting experimental data for these compounds (n > 100,000) is cost-prohibitive, but in-silico approaches^13^ can predict mass spectra for them. For that, three types of approaches exist, rule-based^14^, quantum chemistry^9,10^ and machine learning^13,15^. Additionally, the new deep learning architecture, transformer underling the large-language models (LLMs) which have been driving the recent AI revolution, can be used for predicting in-silico spectra^12^. Large-language models have shown practical applications in biomedical research to summarize clinical notes^16,17^, predict protein structures^18^, estimate binding affinities^19^, and predict biochemical reactions^20,21^.Their capacity to model complex chemical knowledge positions them as a highly valid approach for mass spectra generation.

Here, we generated two comprehensive *in-silico* tandem mass spectral datasets for small molecules, covering the Blood Exposome Database and the Metabolomics Workbench repository. The mass spectral data was segmented into positive ([M+H]+) and negative ([M-H]-) ion modes, encompassing 91,231 of unique endogenous metabolites and exogenous environmental chemicals alongside their structural identifiers and theoretical fragmentation arrays. Comparison analysis showed that our LLM-generated dataset is highly accurate and aligns closely with gold-standard experimental spectra from the NIST23 library. We provide a comprehensive overview of our API-driven data generation pipeline, the structural formatting of the resulting libraries, and extensive technical validation of the predicted mass spectra against gold-standard experimental spectra from the NIST23 database. Our *in-silico* mass spectral datasets could be directly integrated into mass spectrometry data processing pipelines to accelerate high-throughput compound annotation. By providing these datasets as a readily accessible resource, this work aims to expand *in-silico* spectral libraries and accelerate the compound annotation process in mass spectrometry-based metabolomics and exposomics.

## Methods

Pre-trained LLM models can be run locally or accessed by an API offered by commercial providers such as Google or OpenAI. Implementing a local model is resource demanding due to the large number of parameters, so utilizing the online version via an API is preferred for many predictive tasks such as curating a chemical list or summarizing clinical importance of a gene list. Moreover, the Gemini model is available via the online site gemini.google.com and is estimated to be used by over 350 million users per month and is freely available to many academic institutions. This wide accessibility of the Gemini LLM model can democratize the mass spectra prediction and promote a major expansion of mass spectral libraries. We put together a workflow that utilizes the Gemini API to generate mass spectral libraries for small molecules (Figure 1).

**Figure 1:**
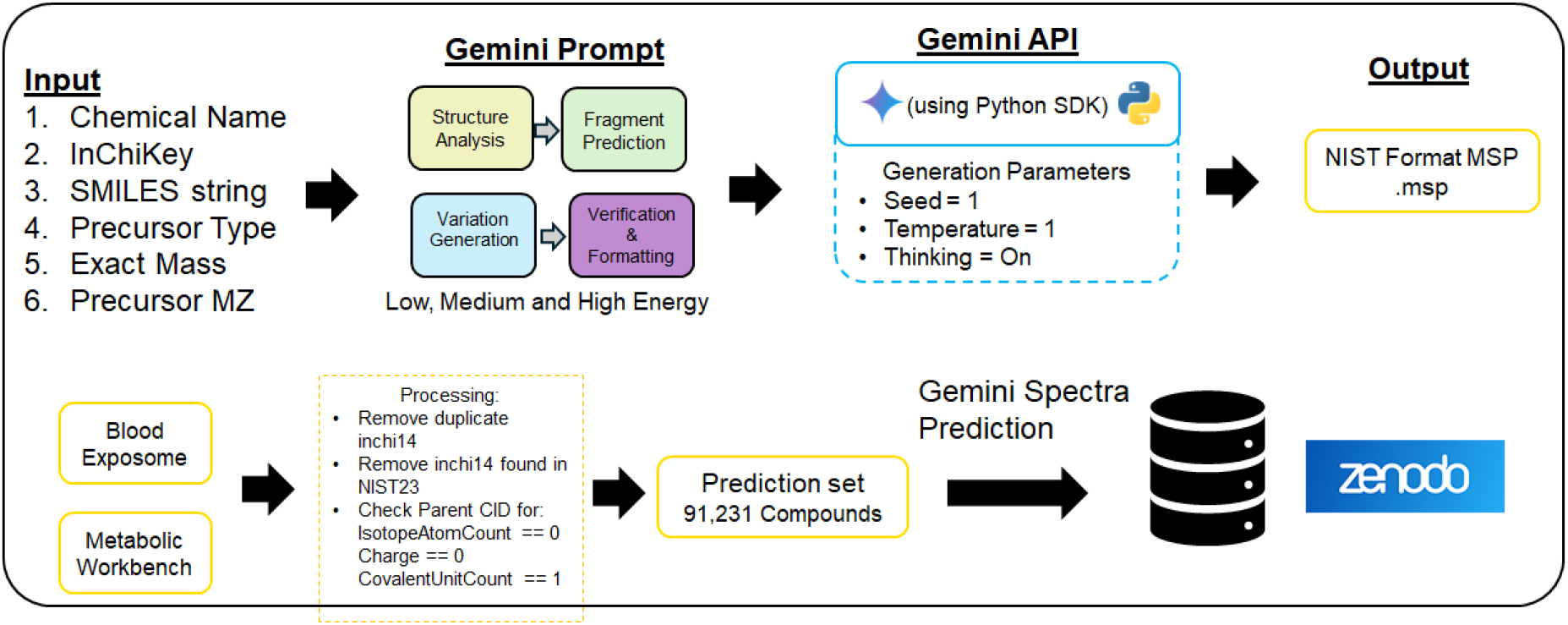
An overview of the mass spectra simulation using the Gemini model.

### Google Gemini Prompt Engineering

We engineered an optimized chain of thought^22^ (CoT) prompts (https://zenodo.org/records/20102592) to instruct Gemini LLM to generate the mass spectra for an input compound. The prompt requires SMILES, Chemical Name, Monoisotopic Mass and Molecular Formula as input, which can be obtained from the PubChem Database^23^. This long-form prompt was designed to guide the model’s reasoning process. This prompt instructed the model to find likely fragmentation mechanisms and stable neutral losses, and then to construct the spectrum based on that reasoning. Examining model’s thinking summary confirms its ability to comprehend and follow detailed guidelines when predicting spectra. We also assessed reproducibility by running predictions for a subset of 100 molecules three times each. The determinism of the model was tested by setting the API’s temperature parameter to zero with fixed seed, which resulted in almost identical outputs for identical inputs, confirming high reproducibility under controlled settings. Prompts were specifically instructed to generate M+H and M-H spectra to cover the electrospray ionization positive and negative modes. The Gemini script was provided with a system-level instruction set prior to receiving any prompts. This instruction defined its role, output format, and constraints. For example, it was instructed to always return the spectrum as a JSON object containing two lists (m/z values and corresponding relative intensities) to ensure consistent, machine-readable output.

### Blood Exposome Database

To test our method on a chemically diverse and biologically relevant set of molecules, we utilized the Blood Exposome Database^1^ (www.bloodexposome.org), also available at https://zenodo.org/records/8146024. This database contains thousands of compounds that have been reported for blood specimens, representing a wide range of chemical classes, including metabolites, environmental contaminants, and food components. The canonical SMILES string for each compound was extracted to serve as the input for our prediction workflow. Chemical properties, including SMILES, Monoisotopic mass and Molecular Formula were collected using PubChem’s PUGREST API (https://pubchem.ncbi.nlm.nih.gov/docs/pug-rest) for each InChiKey.

### Metabolomics Workbench Database

Compounds that were reported in the metabolomics workbench data repository^3^ were downloaded from the PubChem database. It included annotated compounds that were submitted by different depositors as part of their metabolomics studies for different biological organisms.

### Gemini Agent API

All predictions were generated programmatically using the Gemini API. We use the Gemini Pro 3.0 model for the predictions in the batch mode. It took almost three days to simulate spectra for 91,231 unique compounds from the Blood Exposome Database and the Metabolomics Workbench database. A Python script was developed to iterate through the list of unique compounds, which format the appropriate prompt for each, and send requests to the API. The structured spectral data was then parsed from the model’s response and stored for analysis. The final results were exported to the NIST MSP format (Figure 1).

### Validation Against the NIST23 Library

The primary validation was performed by comparing the Gemini-predicted spectra against the gold-standard experimental spectra from the NIST23 library. For each compound, the cosine similarity and entropy similarity score were calculated between the predicted and experimental spectrum using the IDSL.CSA software^5^. The parameters files for the spectra similarity searches are made available at (https://zenodo.org/records/20102592).

### Case study data

We utilized a publicly available study for NIST SRM1950 human plasma reference material. The data is available at https://massive.ucsd.edu/ProteoSAFe/dataset.jsp?task=457c1622e90845d8a004cbe73b105bba. The untargeted DDA data were collected for this study using a Thermo Orbitrap Fusion Lumos mass spectrometer connected with a Vanquish liquid chromatography. Data were processed using IDSL.IPA^24^ and IDSL.CSA^5^ R packages for detecting LCMS peaks and DDA MS2 spectra.

## Data Records

The complete *in-silico* mass spectral datasets generated in this study, alongside the associated prompt parameters, have been deposited in the Zenodo repository and are publicly accessible at https://zenodo.org/records/20102592. The data is released under an open-access Creative Commons Attribution 4.0 International (CC BY 4.0) license to facilitate unhindered use by the metabolomics and exposomics communities.

The data records are organized into two primary files, corresponding to the two distinct chemical databases used in this study 1) Blood_Exposome Database/: This directory contains the simulated tandem mass spectra for the compounds extracted from the Blood Exposome Database. It includes two spectra data files separated by ionization mode – a) blood_exposome_positive_mode.msp: Contains the [M+H]+ adduct spectra. b) blood_exposome_negative_mode.msp: Contains the [M-H]-adduct spectra. 2) Metabolomics_Workbench /: This directory contains the simulated tandem mass spectra for the unique endogenous metabolites and lipids extracted from the NIH Metabolomics Workbench. It includes two files a) metabolomics_workbench_positive_mode.msp: Contains the [M+H]+ adduct spectra. B) metabolomics_workbench_negative_mode.msp: Contains the [M-H]-adduct spectra. Gemini prompts to simulate the spectra are also provided in the Zenodo accession.

These mass spectral libraries are provided in the standard NIST Mass Spectral Format (.msp). The .msp format is a universally recognized, human-readable ASCII text format that is natively supported by virtually all major mass spectrometry data processing software, including MS-DIAL^6^, MZmine^25^, IDSL.CSA^5^, and the NIST MS Search program. Each entry within the .msp file represents a single *in-silico* mass spectrum and is structured as a contiguous text block. The block begins with a standardized set of metadata tags, followed by the spectral peak list (an array of *m/z* and intensity pairs), and is separated from the next entry by a blank line.

To ensure high interoperability and facilitate accurate chemical database searching, each spectral entry contains the following comprehensive metadata fields 1) Name: The common chemical name of the compound, sourced from the PubChem or RefMet databases 2) PrecursorMZ: The exact theoretical mass-to-charge ratio of the precursor ion, calculated based on the designated adduct type (e.g., [M+H]+ or [M-H]-). 3) Precursor_type: The ionization adducts simulated by the LLM (e.g., [M+H]+ or [M-H]-). 4) Formula: The neutral molecular formula of the compound. 5) SMILES: The canonical Simplified Molecular-Input Line-Entry System (SMILES) string. 6) InChIKey: The International Chemical Identifier hash, used for unambiguous structural cross-referencing and database duplication. 7) Num Peaks: An integer indicating the total number of product ions (fragment peaks) predicted by the model and listed in the subsequent array. Following the Num Peaks tag, the *in-silico* peak list is provided. Each line represents a single predicted fragment ion, formatted as two space-separated numerical values: the exact *m/z* of the fragment, followed by its predicted relative intensity. In accordance with standard spectral library conventions (such as those used by MassBank^26^ and NIST), the base peak (the most abundant fragment ion) is normalized to an intensity value of 999, with all other peaks scaled relative to this maximum.

## Technical validation

We performed a series of technical validation and comparisons to show the quality of the generated mass spectral libraries.

We check the overlap of chemical compounds across three major databases: NIST23, Blood Exposome, and Metabolomics WorkBench (Figure 2). A central core of 7,142 compounds is shared among all three databases. However, each database contains a substantial amount of unique data. The Blood Exposome database has the highest number of unique entries at 36,504, closely followed by the Metabolomics WorkBench with 35,669 unique compounds, and NIST23 with 30,662. Pairwise overlaps show that NIST23 shares 4,118 exclusive compounds with Blood Exposome and 5,572 with Metabolomics WorkBench, while the latter two share 5,078 compounds exclusively with each other. It shows that simulated spectra cover many compounds that are expected for blood specimens but are not yet present in the NIST 23 spectra library.

**Figure 2:**
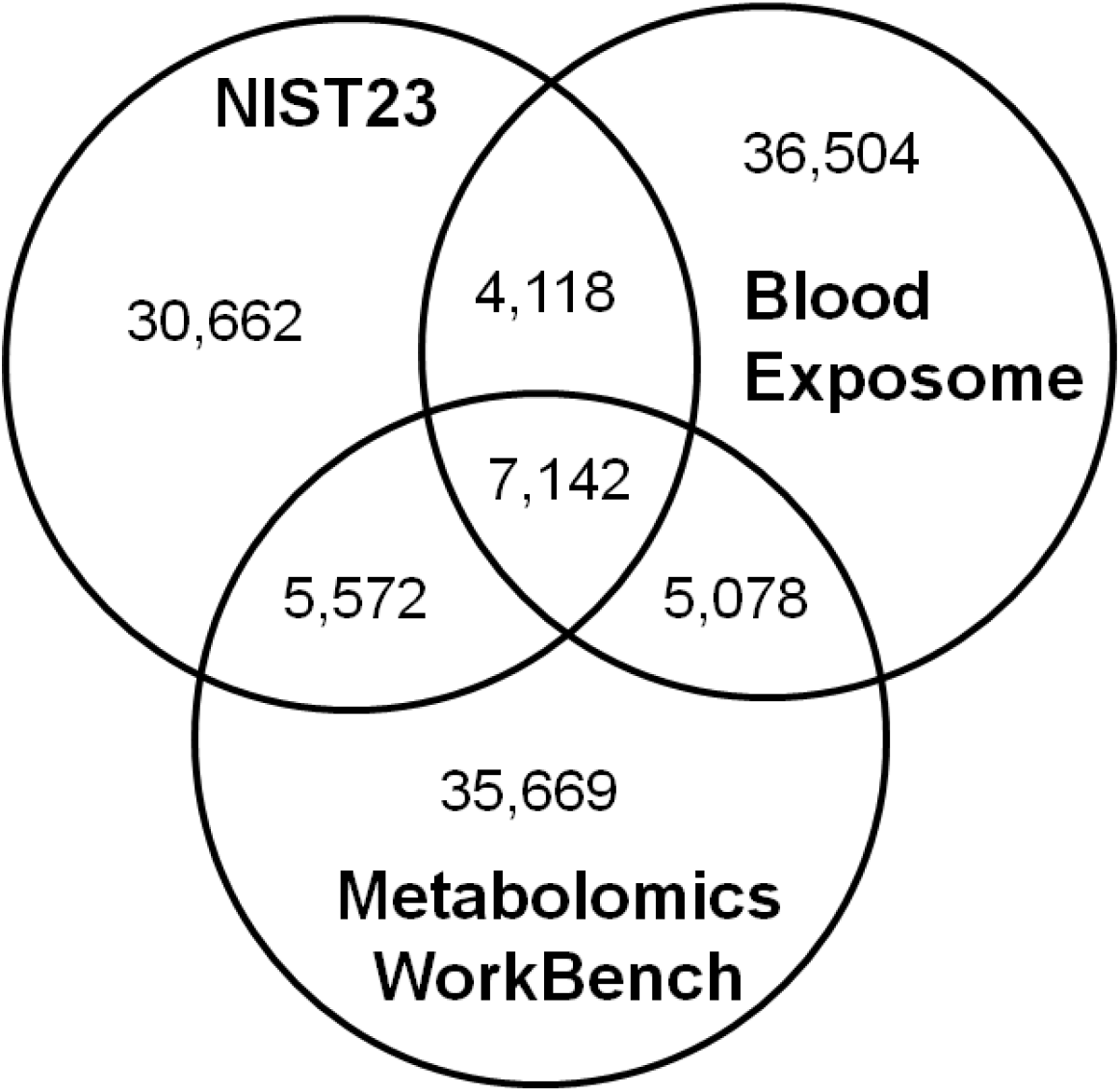
Overlap among major exposomics and metabolomics databases. 2D structure (InChiKey 14) was used for comparison the database contents.

Next, we evaluated the performance of various versions of Gemini models, measuring their ability to generate accurate outputs using two metrics: Mean Entropy Similarity (Figure 3a) and Similarity Hit Count (Figure 3b). Both panels exhibit a consistent trend where advanced “Pro” models and models equipped with “thinking” capabilities significantly outperform earlier or “Lite” versions. The Mean Entropy Similarity peaks with the 3.0 Pro model, achieving the highest score of 0.860. Interestingly, enabling the “thinking” feature in the 3.0 Flash model elevates its score from 0.716 to 0.787, making it highly competitive with the newer 3.1 Pro model (0.7835). Conversely, the 2.5 Flash Lite model performs the poorest, yielding a score of only 0.363. The results were also supported by the spectral similarity hit counts. The 3.0 Pro model leads the group with 782 hits, closely trailed by the 3.1 Pro (723 hits) and the 3.0 Flash (thinking) model (663 hits). The data clearly illustrates an evolutionary improvement across model generations, highlighting that increased model complexity and specific reasoning enhancements yield substantially higher accuracy and fidelity compared to baseline or lightweight iterations. For our work, we have Gemini Pro 3.0 model to simulate the spectra for both blood exposome and metabolomics workbench databases.

**Figure 3.**
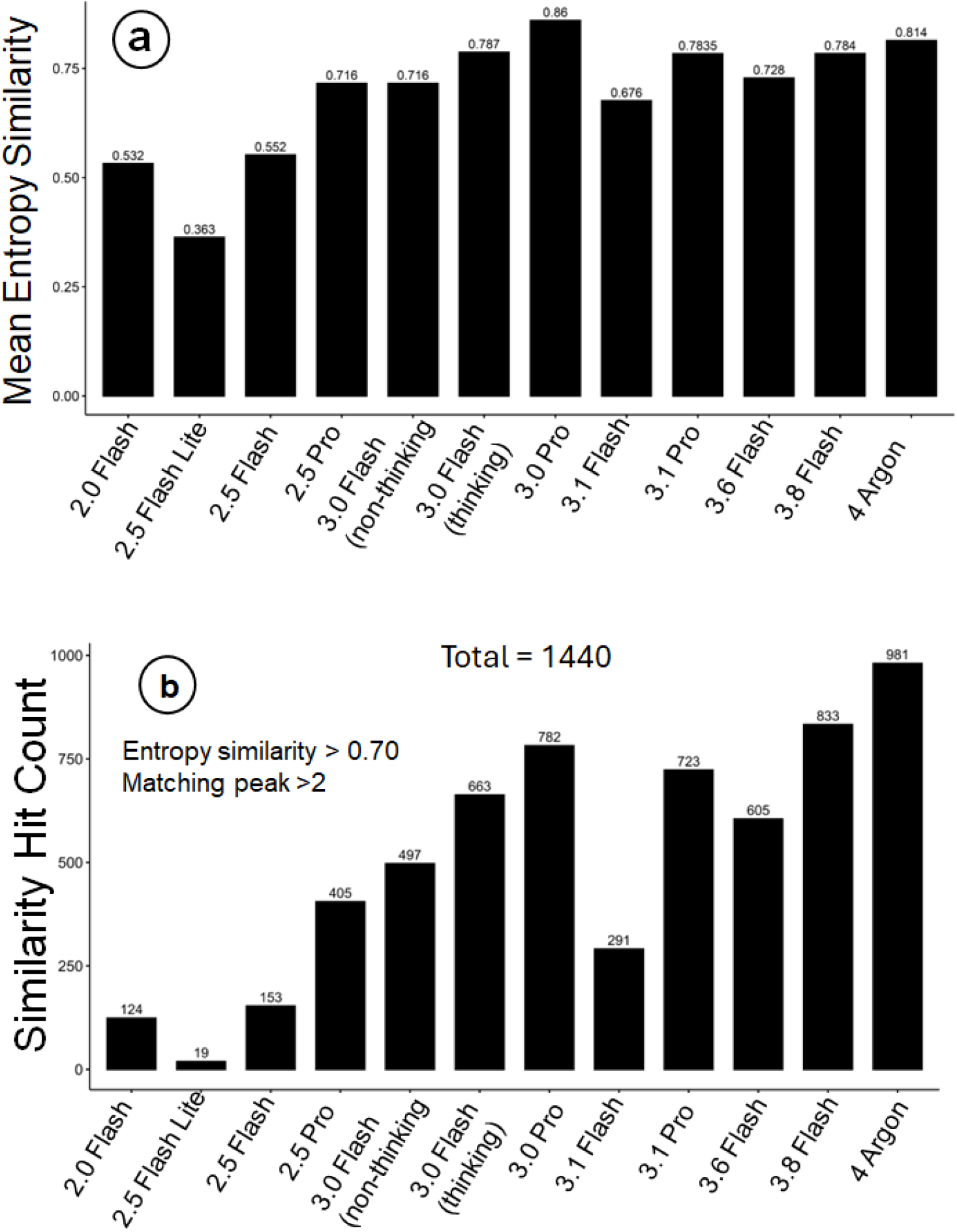
Comparison of the different Gemini Mode versions to simulate spectra for a subset of 1440 compounds that are common among NIST and simulated spectra libraries. a) Median mass spectral entropy similarity between the simulated the NIST entry for the spectra. b) Using the simulated spectra, the annotation for the NIST case study data for NIST SRM1950 samples (see method). Spectral hits above the threshold of entropy similarity 0.70 and at least three matching peaks were considered a hit.

We observed an expected and profound impact of varying collision energies on the fragmentation profiles of small molecules, a critical concept when simulating or evaluating mass spectral libraries (Figure 4). Lower collision energies, specifically 10 eV, result in a constrained number of peaks per spectrum, reflecting minimal molecular dissociation (Figure 4a). As energy increases to 50 eV, the peak count distribution shifts dramatically, yielding highly complex spectra with numerous fragments. It is further supported by an accumulation of low mass-to-charge (m/z) fragments at 50 eV, indicating that the original molecules are aggressively fragmented into much smaller constituents (Figure 4b). Conversely, the 10 eV curve remains dispersed across higher m/z values. Similarly, the intensity of the peaks were also skewed towards lower m/z values at high Collison energy simulations (Figure 4c). These observations validate that the simulated spectral data adheres to expected physical fragmentation rules. Gemini LLM model can logically capture the deterministic reality of energy-dependent molecular breakdown, offering a viable, cost-effective solution to the limited availability of experimental reference libraries.

**Figure 4.**
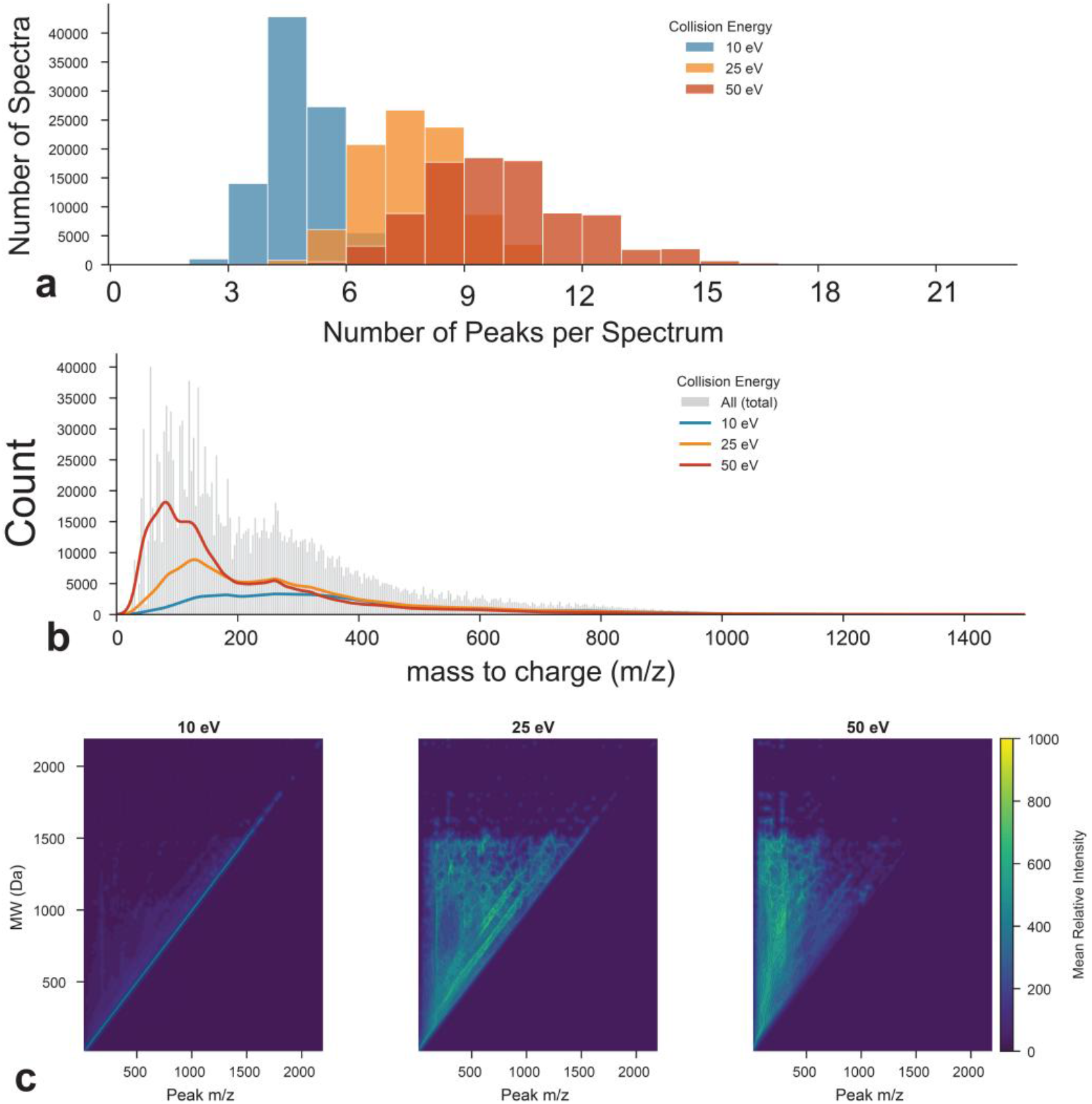
Summary of the spectral data content. a – effect of Collison energy on the peak count. b – effect of Collison energy on the m/z values c) effect of Collison energy on the peak intensity by m/z values.

We observed highly accurate mass spectra simulated by Gemini LLM model for common small molecules. For Acetaminophen, the alignment is exceptionally precise. It correctly returned all the major m/z peaks in the mass spectra and their intensities that are comparable to the true positive spectra in the NIST 2023 database (Figure 5). The model achieved an Entropy Similarity of 0.987 and a Cosine Similarity of 0.998, with major fragment peaks at m/z 110.06 and 152.07 matching nearly perfectly in both presence and relative intensity. Ergothioneine also demonstrates strong predictive concordance, yielding an Entropy Similarity of 0.900 and a Cosine Similarity of 0.931, successfully capturing several key complex fragments, such as those at m/z 127.03 and 186.10. Diazinon exhibits a slightly lower but still robust correlation, with an Entropy Similarity of 0.767 and a Cosine Similarity of 0.922; while the primary base peak at m/z 169.07 aligns well, there is some minor variation in the relative intensities and distribution of smaller product ions. These comparative evaluations demonstrate that general purposed large-language models can successfully predict the high-resolution fragmentation patterns of diverse chemical structures.

**Figure 5.**
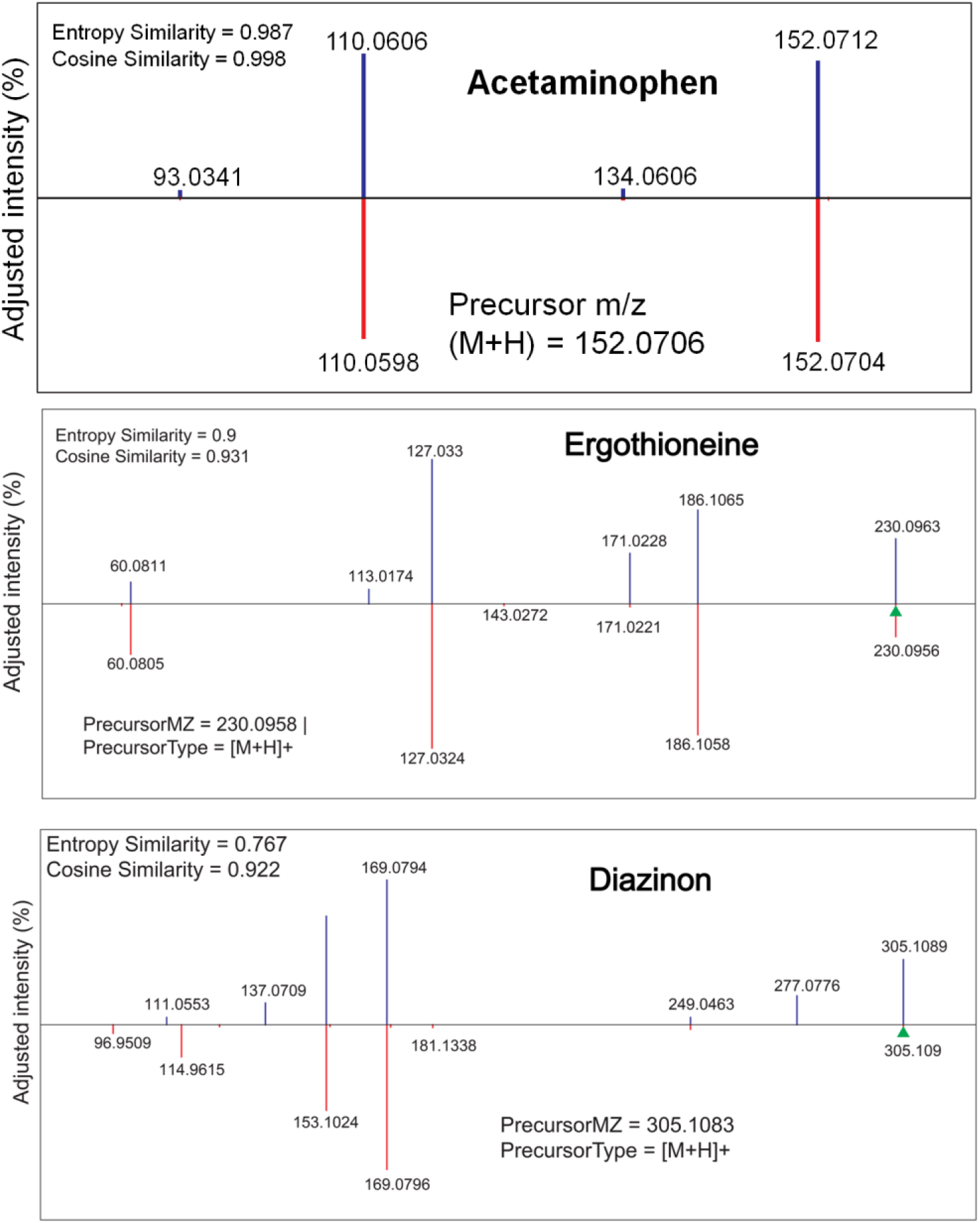
A general-purpose large language model predicted high-quality mass spectra for a small molecule. We used Google’s Gemini LLM model for the prediction of mass spectra. IDSL.CSA software^5^ was used for computing the similarity scores and the draw the spectra similarity plot. These mirror plots compare tandem mass spectra for three distinct small molecules: Acetaminophen, Ergothioneine, and Diazinon. These plots visually align the theoretically predicted mass spectra (top, blue peaks) against standard reference spectra (bottom, red peaks) along a shared mass-to-charge (m/z) axis to evaluate their structural concordance.

The accurate prediction of the mass spectra can be attributed to the mixture of expert and chain-of-thought advancements in the Gemini LLM. The Mixture-of-expert(MoE) architecture^27,28^ which allows LLM to accumulate specialized similar/related domain specific topics knowledge during training by assigning data of similar topics to a subnetwork of the model, called an expert, that develops non-overlapping and focused knowledge based on the assigned topics. It is likely that Gemini training data contains significant amount of material related to mass spectrometry domain, which the MoE architecture and CoT prompt method can take advantage of during spectra prediction.

## Usage Notes

In this Data Descriptor, we present comprehensive *in-silico* tandem mass spectral libraries generated utilizing the general-purpose Google Gemini large language model. Specifically, this descriptor details the programmatic generation of mass spectra for compounds from the Blood Exposome Database, alongside complementary datasets derived from the Metabolomics Workbench. We suggest a practical use of these spectra data in querying experimental DDA data for biospecimens using existing mass spectrometry software. Researchers conducting untargeted liquid chromatography-tandem mass spectrometry (LC-MS/MS) profiling can directly integrate these datasets into their existing data processing pipelines to accelerate the structural annotation of unknown features.

Because the data is formatted in the universal NIST Mass Spectral (.msp) format, it is immediately compatible with the vast majority of open-source and commercial metabolomics software suites. Users can directly import these .msp files into tools such as MS-DIAL^6^, MZmine 3^25^, Global Natural Products Social Molecular Networking^29^, the NIST MS Search GUI, and the IDSL.CSA^5^ package. We recommend using these in-silico libraries as an additional step to searching in-house, NIST23 and existing public mass spectral libraries. We showed an application of the spectral libraries to annotate compounds for the NIST SRM 1950 samples as a case-study. While the Gemini LLM demonstrated high accuracy in replicating established fragmentation rules, it is vital to acknowledge that these are theoretically derived spectra, not empirical ground truths. As such, any structural annotations derived solely from matching experimental data to this Gemini-generated library should be classified as Level 2 (putatively annotated compounds) or Level 3 (putatively characterized compound classes) according to the Metabolomics Standards Initiative (MSI) reporting guidelines. We strongly advise users to employ orthogonal data dimensions to increase the confidence of any annotations driven by this library. Specifically, matching *in-silico* mass spectra should be paired with accurate precursor mass matching (within < 5 ppm mass error), isotopic pattern scoring, and *in-silico* retention time (RT) predictions or collisional cross-section (CCS) matching where applicable. Finally, any features identified using this library that emerge as critical biomarkers or key drivers of biological variance must be strictly validated by purchasing and analyzing a pure synthetic reference standard under identical analytical conditions (MSI Level 1).

## Acknowledgement

The work was in part supported by U24ES035386, R24ES036917, R01ES035478, P30ES023515, UL1TR004419 and R01ES032831.

## Author’s contribution

TN and DB planned the study, implemented the scripts, prepared the results and drafted the manuscript. All authors have reviewed the manuscript’s content. Data availability: https://zenodo.org/records/20102592

